# Riemannian Geometries of Visual Space: Variable Curvature and Horizon

**DOI:** 10.1101/2021.10.25.465820

**Authors:** Jacek Turski

## Abstract

This is a study of the phenomenal geometries constructed in the Riemannian geometry framework from simulated iso-disparity conics in the horizontal visual plane of the binocular system with the asymmetric eyes (AEs). For the eyes’ resting vergence posture, which depends on parameters of the AE, the iso-disparity conics are frontal straight lines in physical space. For all other fixations, the simulated iso-disparity conics consist of families of the ellipses or hyperbolas depending on both the AE’s parameters and the bifoveal fixation. However, the iso-disparity conic’s arcs are perceived in the gaze direction as the frontal lines. Thus, geometries of physical and visual spaces are different. An assumption underlying the relevant architecture of the human visual system is combined with results from simulated iso-disparity straight lines, giving the relative depth as a function of the AE’s parameters and the distance. This establishes the metric tensor in binocular space of fixations for the eyes’ resting vergence posture. The resulting geodesics in the gaze direction, give the distance to the horizon and zero curvature. For all other fixations, only the sign of the curvature may be inferred from the global behavior of the simulated iso-disparity conics.

## 1. Introduction

Visual perception begins at the two eyes when the photons emitted by objects and reflected from surfaces are centrally projected onto retinae and transduced by photoreceptors into electrochemical signals. These signals pass through the multi-layered circuitry of each retina, where they converge on the ganglion cells. The retinal output is sent along the bundle of the ganglion cell’s axons mainly to the primary visual cortex (V1) and continue to more that 30 higher cortices for immensely complex neural processing. However, on their way to V1, axons originating from the nasal half of the retina cross at the optic chiasm to the contralateral brain’s hemisphere and join the temporal half, which remains on the eye’s side of the brain [1]. The splitting and crossing at the optic chiasm underlie our binocular vision [2].

The outcome is the beholder’s three-dimensional visual experience combined with other perceptual modalities and transcended by higher cognitive processes. This article concerns our phenomenal visual space geometric relations before the involvement of psychological processes. To this end, I need first introduce ideas underlying the phenomenon of stereopsis in binocular vision.

When a small retinal area is stimulated in one of the two eyes, there is a unique retinal area in the other eye such that when it is stimulated, a single perceptual phenomenon localized in our phenomenal visual space corresponds to both stimulated areas. These two retinal areas are referred to as corresponding retinal elements and we consider them as having zero disparity. For a given fixation, the horopter is defined as the locus of points in space such that each point on the horopter projects to a pair of corresponding retinal elements. Thus, it is the spatial curve of zero-disparity points.

A small object that is located away from the horopter has nonzero disparity and, in general, is perceived as two objects. However, if the object is located close to the horopter, the brain fuses its disparate images into a single percept and the decussation at the optic chiasm allows images from the two retinae to be compared. The extracted retinal disparity is used by the brain to create our sense of depth relative to the horopter. Even the objects located farther from the horopter, which lead to double vision, convey the sense of depth [3].

More essentially, the difference in retinal disparity between two spatial points is used by the brain to sense the relative depth and, hence, to our perception of objects’ shapes and their relative locations in 3D space, that is, stereopsis and visual space geometry. Because the eyes’ lateral separation, depth perception, visual space geometry and stereopsis are mainly supported by the horizontal retinal correspondence and the spatial longitudinal iso-disparity curves of which the zero-disparity curve is the horopter.

Though the study of phenomenal visual space has had a long history, the geometric relationship between phenomenal visual space and the stimuli-containing physical space remains unresolved. The main reason is that the spatial relations of phenomenal space such as depth and size are, in general, task-dependent and influenced by both contextual factors and our expectations. However, the binocular disparity provides the most precise depth cue that is sufficient for stereopsis with no explicit recognition of a scene’s geometric forms such that the resulting perception is generally unaffected by other factors.

Here, I investigate the disparity-based phenomenal geometry in the binocular system with the asymmetric eyes (AEs) in the framework of the bicentric perspective projections [4]. The AE model is comprised of the fovea’s displacement from the posterior pole by angle *α* = 5.2^*°*^ and the crystalline lens’ tilt by angle *β*. Angle *α* is relatively stable in the human population and angle *β* varies between−0.4^*°*^ and 4.7^*°*^, cf. the discussion in [5]. In the human eye, the fovea’s displacement from the posterior pole and the cornea’s asphericity contribute to optical aberrations that the lens’ tilt then partially compensates for [6].

The horopteric conics simulated in [4] resemble the empirical horopters and are extended in this study to the families of iso-disparity conics for a stationary, upright head with the eyes converging on the points of the horizontal binocular field of fixations. For the eyes in the resting vergence posture, that is, when the eyes fixate at the abathic distance, the iso-disparity curves are straight frontal lines. For other fixations, the iso-disparity curves consist of either a family of ellipses or hyperbolas, depending on the AE’s asymmetry parameters and the location of the fixation point.

However, the iso-disparity curves are always perceived in the gaze direction as the frontal lines. Therefore, the geometries of physical space and phenomenal visual space are different and the global aspects of this difference are studied here in the framework of Riemannian geometry. The appendix introduces the most basic concepts of Riemannian geometry in one global chart, the setting of this study that is imposed by a stationary, upright head.

## 2. Retinal Correspondence

The correspondence of retinal elements, as it is described in the first section, forms the basis of the nonius paradigm for a direct determination of the corresponding retinal elements [7, 8]. All other methods, as reviewed in [9], are indirect and less reliable.

However, retinal correspondence is not readily available in nonius measurements of longitudinal horopter [9]. More importantly, it cannot be obtained from geometrical methods because of the beholder eye’s asymmetry: the corresponding points are compressed in the temporal retinae relative to those in the nasal retinae [10]. This situation changed when the geometric theory of the binocular system with AEs was developed in [4] that fully specified the retinal correspondence in terms of AE’s asymmetry parameters. It was verified in simulations of the horopteric conics starting from the resting vergence eyes’ posture shown in Figure 1.

**Figure 1:**
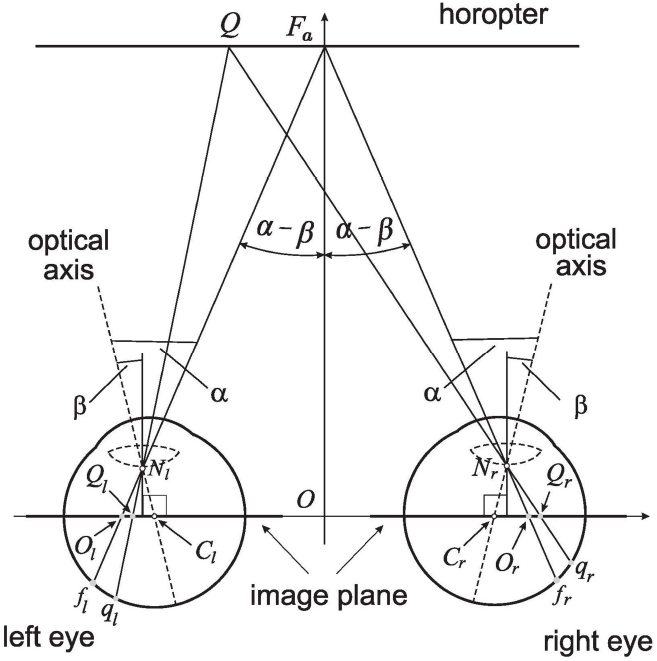
Retinal correspondence. The linear horopter passing through the fixation *F*_*a*_ at the abathic distance *d*_*a*_ =|*OF*_*a*_|. The fixation point *F*_*a*_ projects to the foveae *f*_*r*_ and *f*_*l*_ and the point Q on the horopter projects to the retinal corresponding points *q*_*r*_ and *q*_*l*_ and to points *Q*_*r*_ and *Q*_*l*_ in the image plane. The subtense *σ*_*a*_ at *F*_*a*_ is given by 2(*α* − *β*) [4].

I briefly review the retinal correspondence in the binocular system with AEs established in [4] because in Section 4 it is used to formulate the extension of the horopteric conics (zero-disparity conics) to the iso-disparity conics.

The distribution of the retinal corresponding points *q*_*r*_, *q*_*l*_ is asymmetrical with respect to the foveae *f*_*r*_ and and *f*_*l*_ such that |*q*_*r*_*f*_*r*_| = |*q*_*l*_*f*_*l*_|, see Figure 1. It is demonstrated in [4], that the distribution of these corresponding points projected into the image planes of each AE, *Q*_*r*_, *Q*_*l*_, satisfy |*Q*_*r*_*O*_*r*_| = |*Q*_*l*_*O*_*l*_| where *O*_*r*_ and *O*_*l*_ are projected foveae *f*_*r*_ and *f*_*l*_, respectively. It is proved in [4] that this symmetry on the image plane is preserved for all other fixations. Thus, unknown (asymmetric) retinal correspondence can be entirely formulated on the image plane of the AE in terms of the asymmetry parameters.

The abathic distance *d*_*a*_ = |*OF*_*a*_| to the fixation point *F*_*a*_ in the eyes’ resting vergence posture was obtained in [4] as follows:

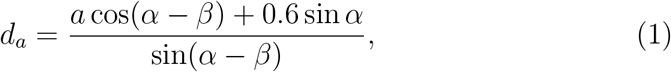

where 2*a* = 6.5 cm is the ocular separation and 0.6 cm is the distance of the nodal point from the eyeball’s rotation center.

I close this section by explaining why the eyes fixating at the abathic distance are referred to as the eyes’ resting vergence posture, or later as the resting eyes for short. The image plane in the AE model is parallel to the lens’s equatorial plane and is passing through the eye’s rotation center. Figure 1 shows the eyes’ posture in which the image planes are coplanar and the horopter is frontal straight line passing through the fixation point at the abathic distance. This distance numerically corresponds to the eyes’ resting vergence posture [11]. Because the eye muscles’ natural tonus resting position serves as a zero-reference level for convergence effort [12], this eye posture is identified in [4] with the eyes’ binocular primary position.

## 3. Binocular Field of Fixations

For an observer with a stationary, upright head and binocular fixations in the horizontal plane, the coordinate system consists of the *y*-axis passing through the eyes’ rotation centers and the head *z*-axis formed by the intersection of the midsagittal plane with the horizontal visual plane, see Figure 2.

**Figure 2:**
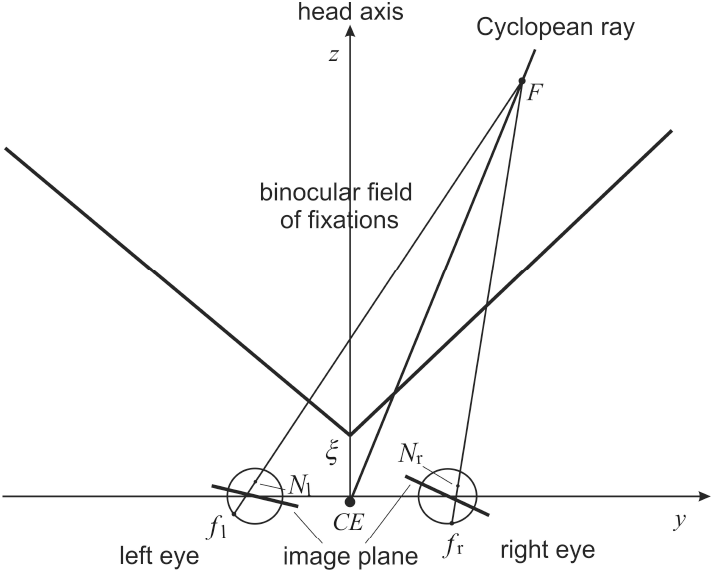
The binocular field of fixations in the horizontal visual plane for the binocular system with the AEs.

The binocular field of fixations is limited by the head/eyes anatomy and physiology: the protruded nose and the fixations neurally restricted to the azimuthal range ± 45^*°*^ [13]. Further, the geometry of the eyes in symmetric convergence shown in Figure 1, demonstrate that the binocular system possesses bilateral symmetry—the symmetry with respect to the head’s midsagittal plane. Therefore, the binocular field of fixation, denoted by *B*, is assumed symmetric about the z-axis. Thus, *B* is a convex set shown in Figure 2 where the point (0, *ξ*) denotes the closest point on which the eyes can converge. For fixation point, the monocular perception from outside of *B* is still available. In fact, the perceived Cyclopean direction, specified by the Cyclopean ray in Figure 2, does not depends on whether the stimulus reaches the retinal element in one eye or its corresponding element in the other eye alone or whether it reaches both the corresponding element simultaneously [9]. However, stereopsis is not available in the region outside of *B*.

For a fixation point, the local chart for the binocular visual space is the whole set *B*. For every fixation (*y, z*)∈ *B*, I want to find the corresponding scalar product on the tangent plane, *R*^2^, with its origin at (*y, z*), using the biologically motivated geometric theory of human binocular perception developed in [4]. These families of scalar products will define the Riemannian geometries of visual space, each dependent on both the AEs parameters and the point of binocular fixation. Underlying notions of Riemannian geometry are outlined in Appendix Appendix A.

## 4. Simulation of Iso-Disparity Curves

The iso-disparity conics are first specified on the image planes of the AEs when the eyes are in the resting vergence posture shown in Figure 1. Referring to this figure, recall that points *Q*_*r*_ and *Q*_*l*_ on the same side and of the same distance from the centers *O*_*r*_ and *O*_*l*_ are the corresponding points covering the corresponding retinal elements *q*_*r*_ and *q*_*l*_. The corresponding points back-projected through the nodal points give the spatial point *Q* that is on the horopter line.

The iso-disparity lines are constructed as follows. The points *Q*_*r*_ + *δ*_0_ and *Q*_*l*_ on the *y*-coordinate line, when back-projected, gives the crossed point on the *δ*_0_-disparity line. Similarly, the points *Q*_*r*_ − *δ*_0_ and *Q*_*l*_, when back-projected, give the uncrossed point on the −*δ*_0_-disparity line.

The simulations of the iso-disparity conics of a constant disparity difference that are carried out in *GeoGebra* are shown in Figure 3, Figure 4, and Figure 5 for the same relative disparity *δ*_0_ = 0.00355 cm.

**Figure 3:**
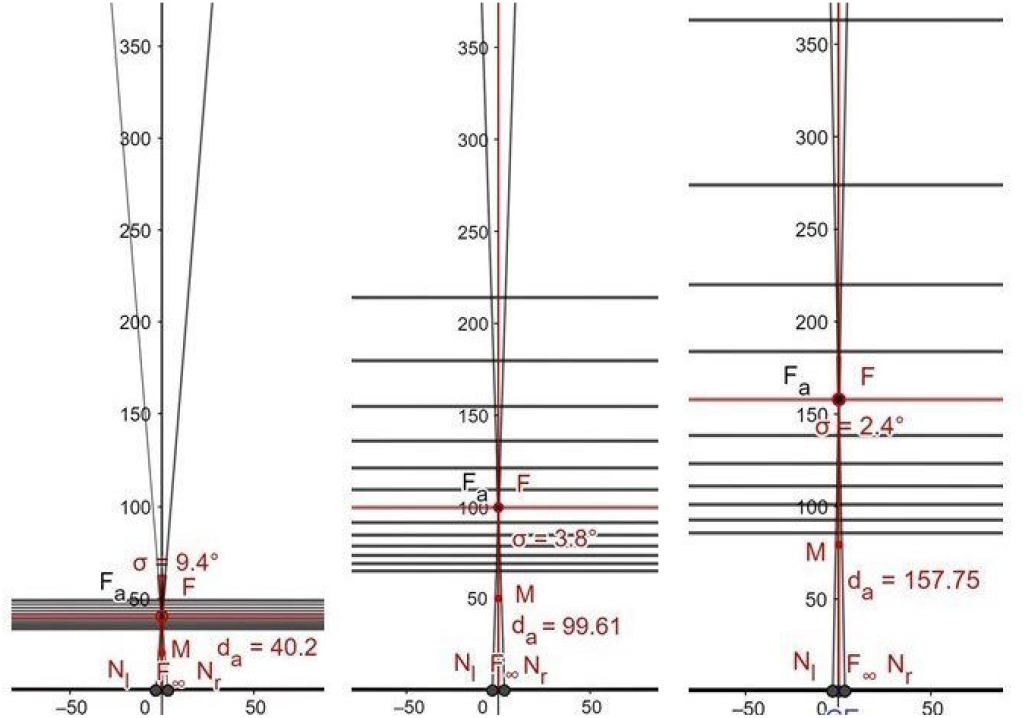
The iso-disparity lines for the eyes’ resting vergence fixations with the abathic distance given in (1): on the left, the abathic distance *d*_*a*_ = 40.2 cm for *β* = 0.5^*°*^, in the middle, the abathic distance *d*_*a*_ = 99.61 cm for *β* = 3.3^*°*^, and on the left, the abathic distance *d*_*a*_ = 157, 75 cm for *β* = 4^*°*^. The angles at *F* are *σ* = 2(*α*− *β*), where *α* = 5.2^*°*^. The horopters are shown in horizontal red lines.

**Figure 4:**
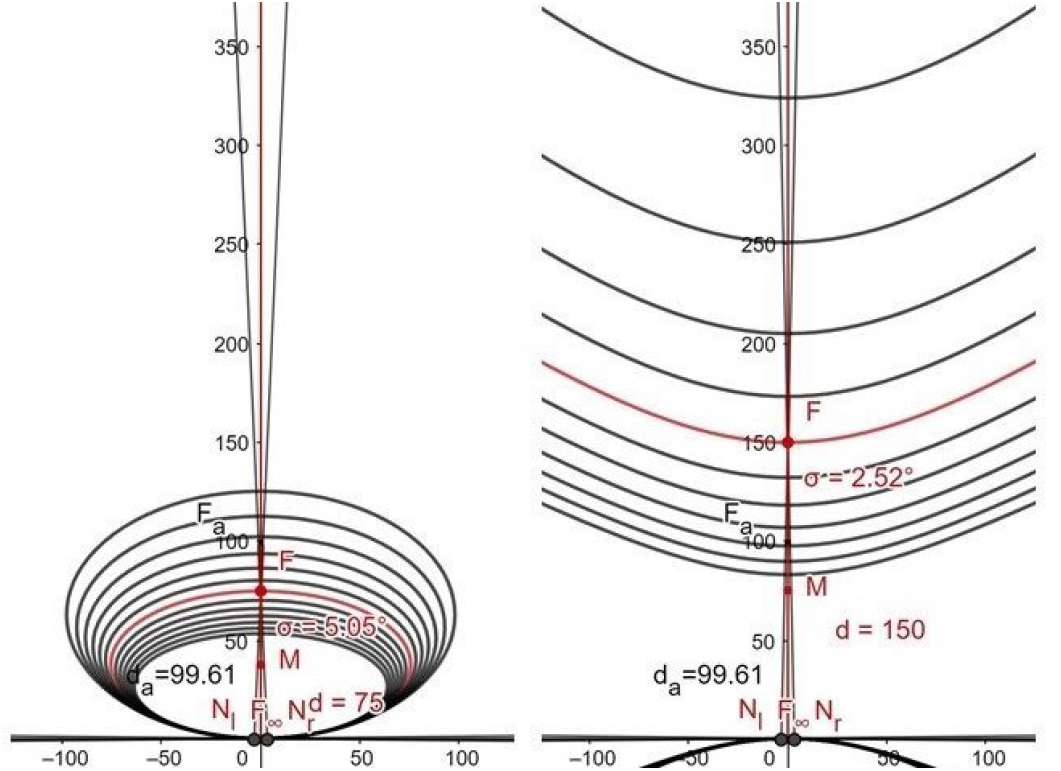
The iso-disparity conics for the binocular system of the abathic distance *d*_*a*_ = 99.61 cm for two symmetric fixations at the indicated distances *d*: ellipses on the left and hyperbolas on the right. The conics in red lines are horopters.

**Figure 5:**
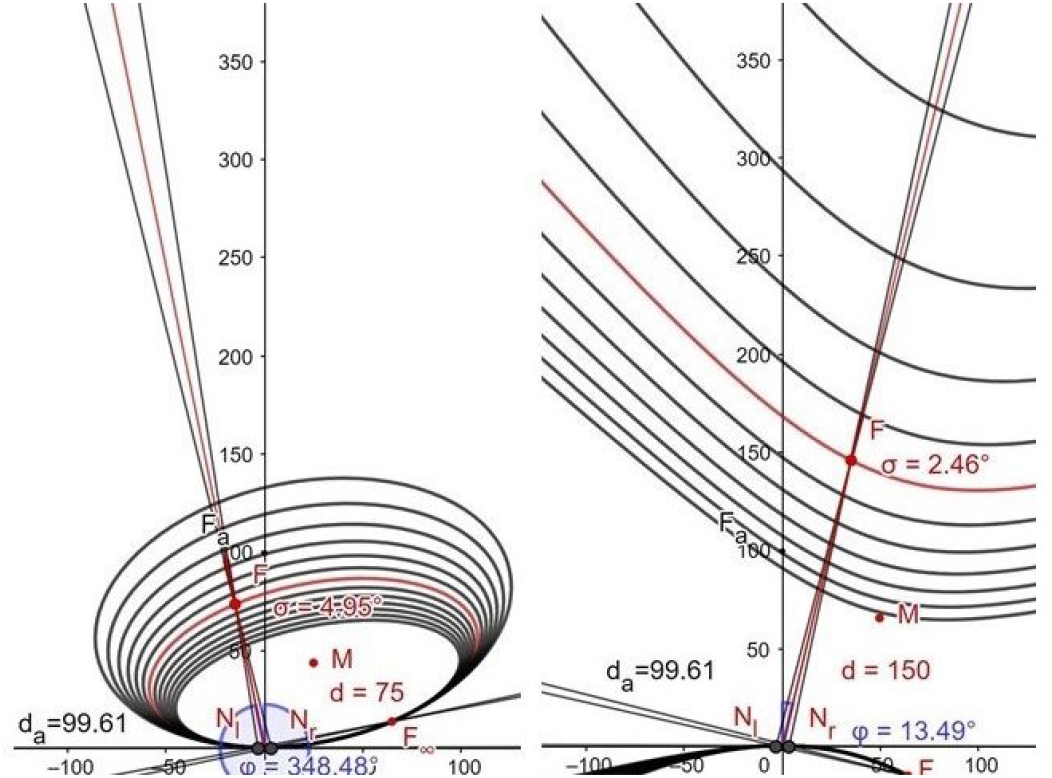
The iso-disparity conics for two asymmetric fixations: ellipses on the left and hyperbolas on the right. The conics in red lines are horopters. The straight line passing through the fixation *F* is the Cyclopean ray. Its direction specifies the orientation of the conics. The indicated distances *d* are measured along the Cyclopean ray.

Figure 4 and Figure 5 show how the iso-disparity lines for the case of *d*_*a*_ = 99.61 cm in Figure 3 are transformed into ellipses and hyperbolas for symmetrically (Figure 4) and asymmetrically (Figure 5) convergent eyes in the binocular field. The fixation points are of the distance 75 cm and 150 cm along the Cyclopean ray for the ellipses and hyperbolas, respectively. The direction of the Cyclopean ray specifies the orientation of the horopteric conics [4] and, therefore, the orientations of the iso-disparity conics for a given fixation.

## 5. Relative Depth for Resting Eyes

From the simulations shown in Figure 3 for the disparity step of *δ*_0_ = 0.00355 cm, I calculate the distance Δ*z*_*i*_ and midpoint *z*_*i*_ between the consecutive iso-disparity lines to approximate how the depth of iso-disparity lines changes with distance to the head in the gaze’s direction. The points (*z*_*i*_, Δ*z*_*i*_) for different abathic distance eyes fixations lie on the graph of a single quadratic function

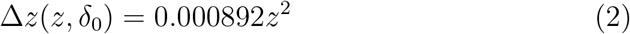

shown in Figure 6.

**Figure 6:**
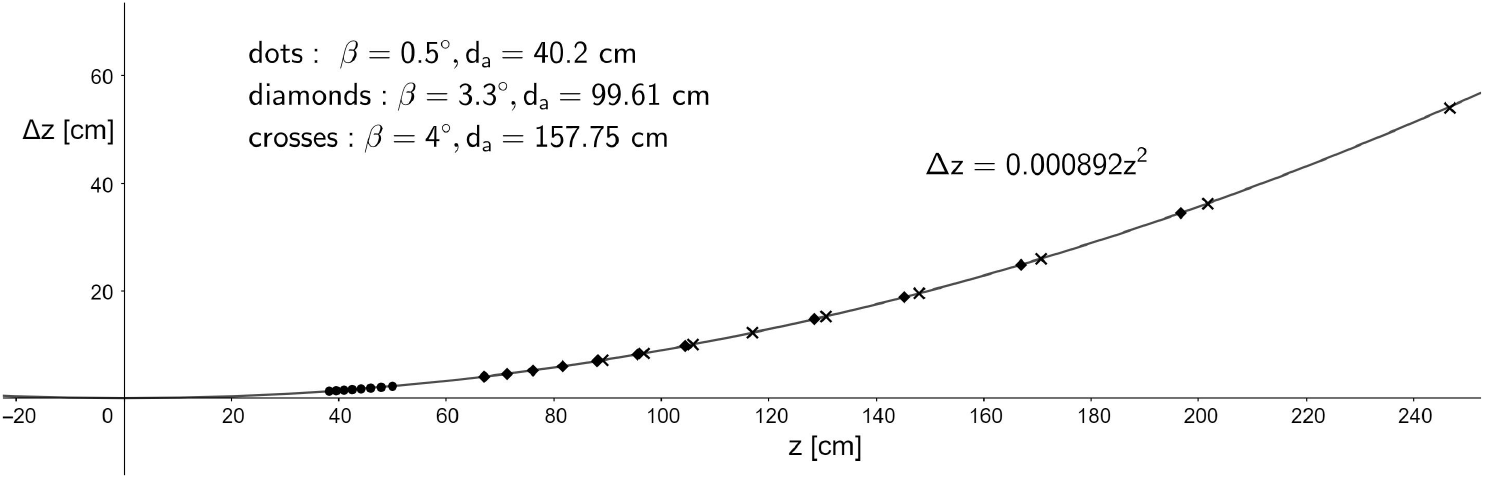
For the simulations shown in Figure 3, the points (*z*, Δ*z*) are plotted for three different abathic distance fixations: *d*_*a*_ = 40.2 cm (dots), *d*_*a*_ = 99.61 cm (diamonds), and *d*_*a*_ = 157.75 cm (crosses).

To investigate the iso-disparity lines distribution dependence on disparity value, I compare simulated iso-disparity lines for disparity difference *δ*_0_ = 0.00355 cm with lines simulated for disparity *δ* = (1*/*5)*δ*_0_ = 0.00071 cm, each for the abathic distance *d*_*a*_ = 99.61 cm. The result of these two simulations is shown in Figure 7.

**Figure 7:**
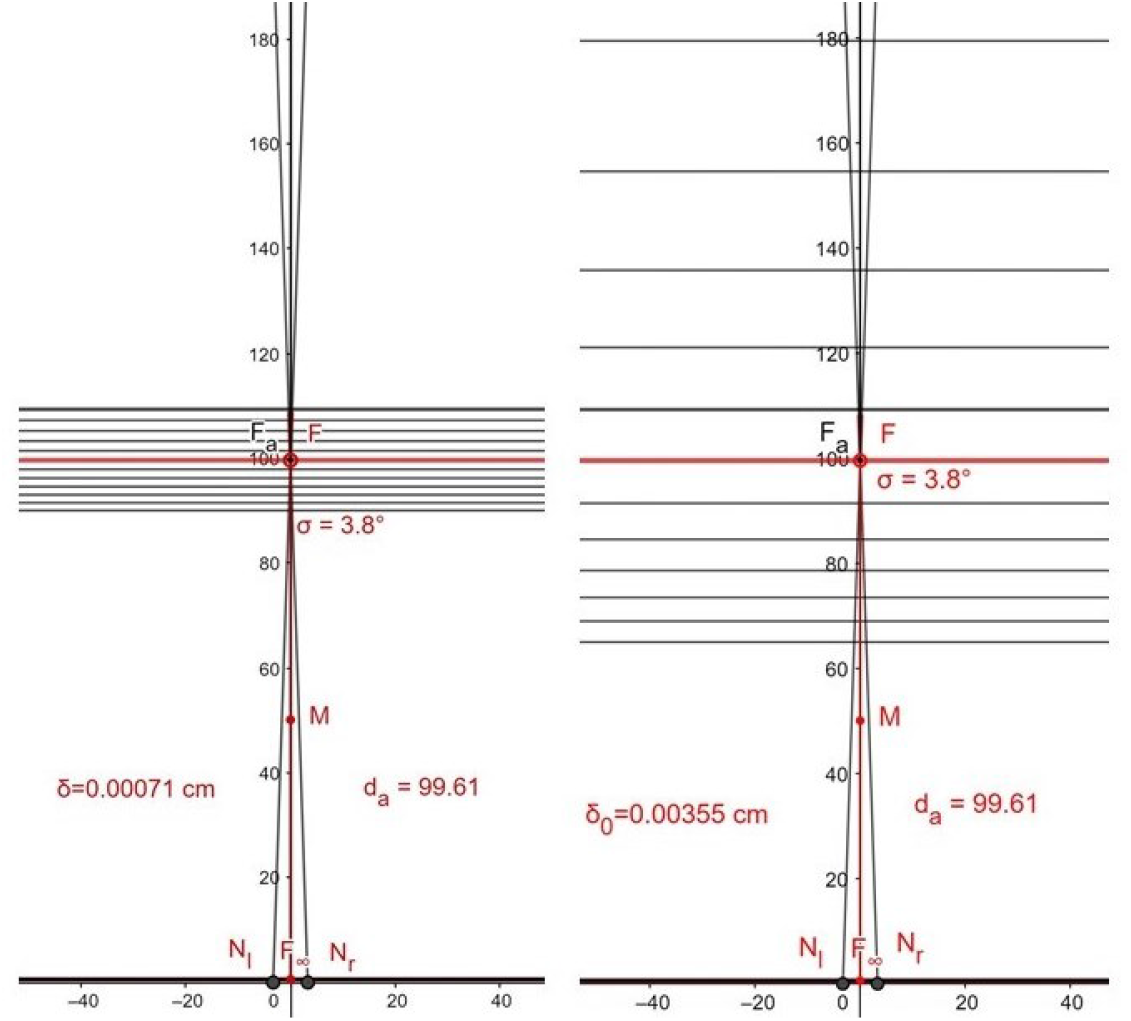
The is-disparity lines for *d*_*a*_ = 99.61 cm with different disparity values; on the right, *δ*_0_ = 0.00355 cm, while on the left, *δ* = 0.00071 cm.

We can see that each space between the neighboring iso-disparity lines for disparity *δ*_0_ = 0.00355 shown in Figure 7’s right panel is filled by an additional five iso-disparity lines for disparity *δ* = 0.00071 shown on the left in Figure 7. Thus, the relation

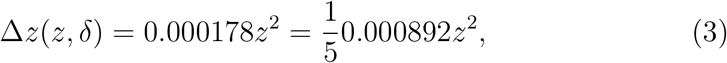

can be predicted from the simulations and verified numerically. In fact, substituting *d*_*a*_ = 99.61 cm for *z* in (3), we obtain,

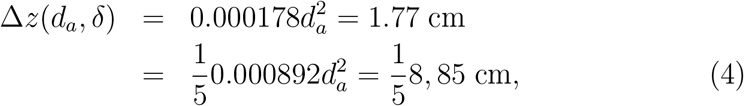

which agree with the simulations shown in Figure 7.

The above discussion of simulations demonstrate that for any *z >* 0, disparity *δ*, and *h >* 0,

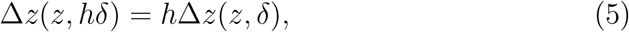

This relation can be scaled down to the value of the disparity imposed by the foveal distribution of photoreceptors.

I conclude that the interval between iso-disparity lines for the disparity difference *δ* = *hδ*_0_ satisfies the relation,

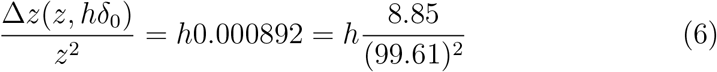

where 8.85 cm is the distance between the horopter line and the −*δ*_0_-disparity line when *d*_*a*_ = 99.61 cm for *β* = 3.3^*°*^.

I choose the *h* value such that Δ*z*(*d*_*a*_, *hδ*_0_) gives the depth resolution at the fovea where the cone-to-cone spacing is about 2.7×10^*−*4^ cm, which gives the corresponding spacing of 10^*−*4^ cm on the image plane of the AE. Given that *δ*_0_ = 0.00355 cm, we obtain *h* = 10^*−*4^*/*3.55×10^*−*3^≈1*/*35. Thus, for *h* = 1*/*35,

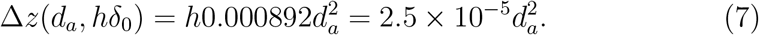

To simplify notation, I denote Δ*z*(*z*, (1*/*35)*δ*_0_) by Δ*z*(*z*). Then, we obtain from (7): Δ*z*(*d*_*a*_) ≈ 0.25 cm for *d*_*a*_ = 99.61 and Δ*z*(*d*_*a*_) ≈0.04 cm for *d*_*a*_ = 40.2.

In order to model relative depth, the simulations must be combined with the fundamental features of the human visual system’s architecture, which include the cones’ eccentricity dependent density and the convergence of the photoreceptors on the ganglion cells, which is 1-to-1 in the foveala and which approaches 200-to-1 in the periphery. Related to this issue, the concept of receptive fields becomes important. Visual information is sent from the eyes along the ganglion cells’ axons and arrives, mainly, at the primary visual cortex, where it is retinotopically mapped with a significant magnification of the foveal region and scaled logarithmically with retinal eccentricity [14].

There is no theory based on first principles that can describe how perceived relative depth varies with disparity and distance. Therefore, using (6), (7) and the above discussion, I postulate that the relative depth element *δp* in visual space approximately satisfies the relation.

### Postulate.

*The relative depth element δd in visual space is given by*

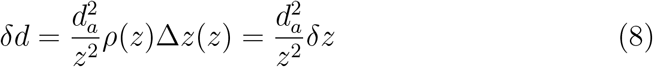

*where δz* = Δ*z*(*d*_*a*_). *Here ρ*(*z*) *accounts for the anatomical and physiological aspects of the human visual system*.

Human disparity contains two components, fine disparity and coarse disparity [15]. Fine disparity allows us to determine the depth of objects in the central visual area where disparate images are fused into a single percept (Panum’s fusional area). Coarse disparity provides stereopsis from disparities well beyond the fusible range. It is still useful for depth perception for moderately large disparities even for double images but is only clearly signed with a vague impression of depth magnitude for very large disparity values [16, 17]. Importantly, coarse disparity creates our sense of being immersed in the ambient environment [18].

I want to emphasize in this work the modeling of the global aspects of the visual space geometry that can be studied in the Riemannian geometry framework. Some of the global aspects of visual space geometry, which seem to agree with the experimental results reported in the literature, are uncovered by the theory developed here from the relative depth formula postulated in (8).

## 6. Metric Tensor for Resting Eyes

In this section, I propose a model of the visual space metric tensor, the fundamental concept of Riemannian geometry. To do this, I first derive the expression for the line element in the *y*-direction.

The length of *δy* element on the iso-disparity line passing through (0, *z*_1_ and projected to the Cyclopean eye image is directly proportional to the length of line element *δy* and inversely proportional to the distance to the iso-disparity line. Assuming the veridical line element perception *δλ* at the abathic distance,

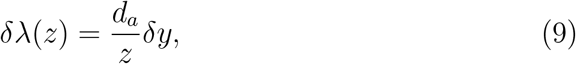

and combining it with (8), we obtain the visual space metric line element

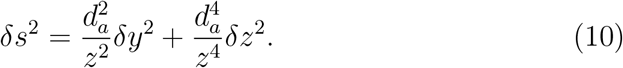

The binocular field of fixations *B* in the horizontal plane was discussed in Section 3. This is a convex set parametrized by the coordinates (*y, z*) that are shown in Figure 2. The set *B* has the Cyclopean visual metric determined by a map

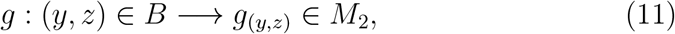

where *g*_(*y,z*)_ is the metric tensor obtained from (10),

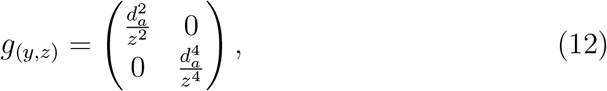

and *M*_2_ is the set of all real symmetric positive defined 2×2 matrices.

The visual metric tensor defines an inner product on each tangent plane *T*_(*y,z*)_*B*, the tangent plane *R*^2^ at (*y, z*) that has the origin at this point. Because (0, 0) ∉ *B*, the map *g* in (11) is a smooth function on *B*. The inner product is usually written⟨*∂*_*j*_, *∂*_*k*_⟩= *g*_*p*_(*∂*_*j*_, *∂*_*k*_) = *g*_*jk*_(*p*) where *p* = (*y, z*) and each index runs over *y* and *z* for the vector basis *∂*_*y*_, *∂*_*z*_ ∈ *T*_(*y,z*)_*B*.

Although I use standard Riemannian geometry notation for writing the basis vectors, cf. Appendix, it should be understood that for the coordinates (*y, z*), *∂*_*y*_ = (1, 0), and *∂*_*z*_ = (0, 1). Thus, the length of the vector *V* = (*V* ^*i*^), is 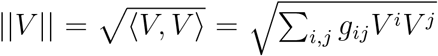 and the angle between vectors *U* and *V* can be obtained from cos *θ* =⟨*U, V*⟩*/*||*U*||||*V*||.

The pair (*B, g*) is a global chart on the binocular field that allows computing length of vectors in tangent spaces and, therefore, length of curves in *B*. It emphasizes global aspects of the visual space. For the more precise notion of the tangent vector field on a manifold that is covered by only one chart, I refer to Appendix Appendix A.1.

## 7. Geodesic Equations for Resting Eyes

For an excellent exposition of the Riemannian geometry framework, I refer to [19]. A short introduction to the notions of covariant derivative, connection and parallel transport, which underlie the fundamental concept of geodesics, is given in Appendix Appendix A.2 and Appendix A.3.

I start by writing the defining expressions for the Christoffel symbols

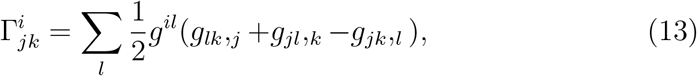

where (*g*^*il*^) = (*g*_*il*_)^*−*1^ is the inverse matrix.

Then, a geodesic is a curve *γ*(*t*) = (*x*^*i*^(*t*)) whose the covariant derivative of the velocity vector is zero,

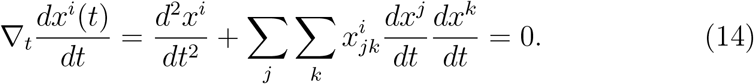

In the notation used for coordinates (*y, z*), the tensor matrix (12) entries

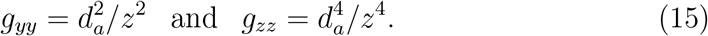

The non-zero Christoffel symbols (13) are the following:

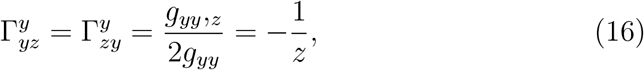

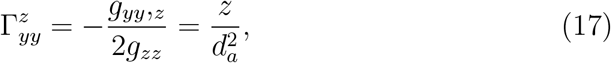

and

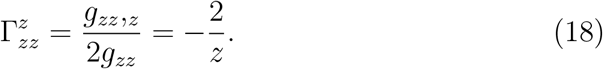

Thus, in the notation of this paper, I obtain the differential equations for the geodesic *γ*(*t*) = (*y*(*t*), *z*(*t*) as follows.

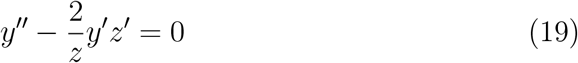

and

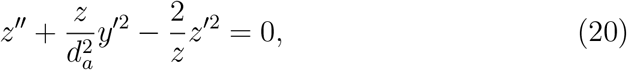

where the ‘prime’ means ‘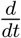’. I consider two sets of the initial conditions,

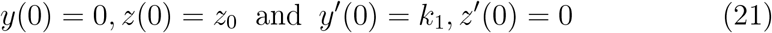

and

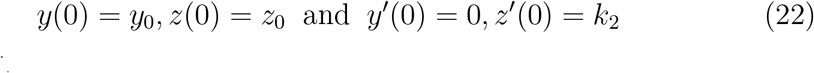

where *z*_0_ *> ξ*.

To solve this system of equations with the first set of initial conditions (21), I start by integrating (19) to obtain 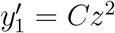. Here, subscript ‘1’ indicates the solution for the initial conditions (21). Later, subscript ‘2’ will be used for the solution of the second initial conditions (22). By substituting the solution for *y*^*′*^ into (20) and rewriting the equations in terms of (*y*_1_(*t*), *w*_1_(*t*)) where *w*_1_(*t*) = 1*/z*_1_(*t*), we obtain the system of geodesic equations

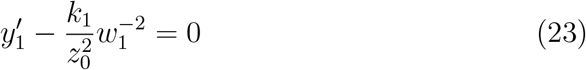

and

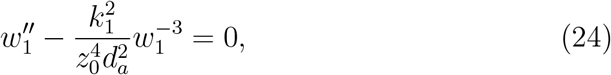

satisfying the initial conditions (21).

The equation (24) is the simplest of the celebrated Ermakov-Pinney equations [20, 21] distinguished be the fact that while being nonlinear, they have general solutions. Following [21], for the initial conditions *w*(0) = 1*/z*_1_, *w*^*′*^(0) = 0, the solution of (24) satisfying the initial conditions is

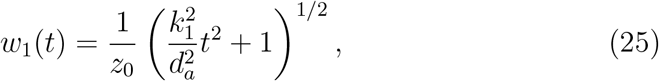

For the second set of initial conditions (22), we have 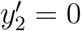 and, then,

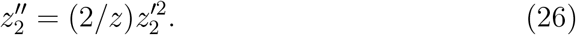

The geodesics for both initial conditions are derived in the next section where also their basic properties are analysed.

## 8. Geodesics in Resting Eyes Visual Space

For the first set of initial conditions, substituting (25) into (23), we derive the expression for 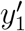

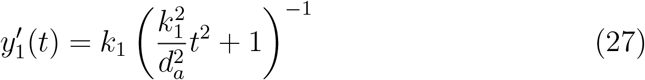

which can be easily integrated,

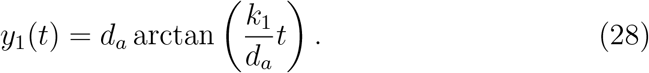

Before I derive the geodesics, I need to calculate *z*^*′*^(*t*). Rewriting (25) as *z*(*t*) = 1*/w*(*t*) and differentiating it, we obtain,

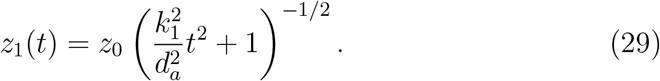

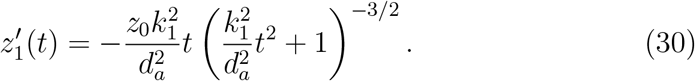

For the second set of the initial conditions, integrating (26), we first have

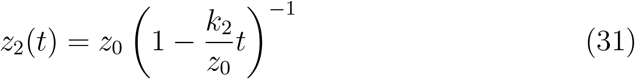

and then we obtain *y*_2_(*t*) = *y*_0_ and

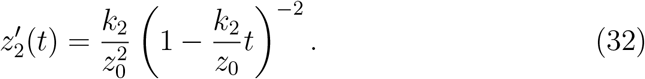

### Theorem.

*In the resting eyes visual space there are two families of parallel geodesics. The first type of geodesics are starting at a point* (0, *z*_0_) ∈ *B in the direction ∂*_*y*_ *and the other type of the geodesics are starting at* (*y*_0_, *ξ*) *in the direction ∂*_*z*_. *Here ξ is the closest point on the z-axis on which the eyes still can converge*.

*Proof*. I prove this theorem by showing that the curves satisfying the geodesic equations first starting at (0, *z*_0_) in the directions of *∂*_*y*_ and then starting at (*y*_0_, *ξ*) in the direction of *∂*_*z*_ have unit speed parametrizations for the corresponding values of *k*_1_ and *k*_2_, respectively. For the direction vector *∂*_*y*_, I solve the unit speed condition for the curve *γ*_1_(*t*) = (*y*_1_(*t*), *z*_1_(*t*))

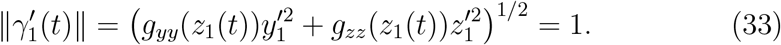

I obtain the unit speed condition

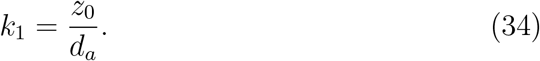

Therefore, in this case, the geodesic is given by

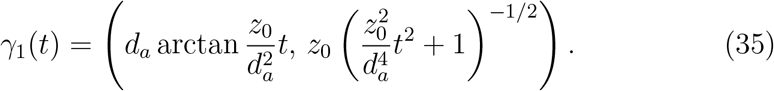

For the direction vector *∂*_*z*_, the unit speed condition

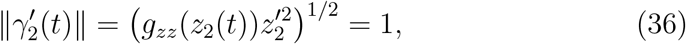

gives

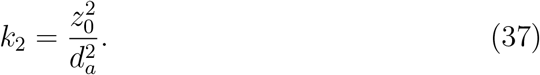

The corresponding geodesic has the expression

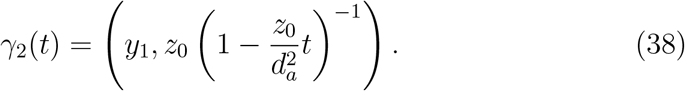

The graphs of the geodesics for the abathic distances *d*_*a*_ = 40.2 cm, *d*_*a*_ = 99.61 cm and *d*_*a*_ = 157.75 cm shown in Figure 8 demonstrate the the geodesics for a given *d*_*a*_ value in each of family do not intersect, that is, they are parallel.

**Figure 8:**
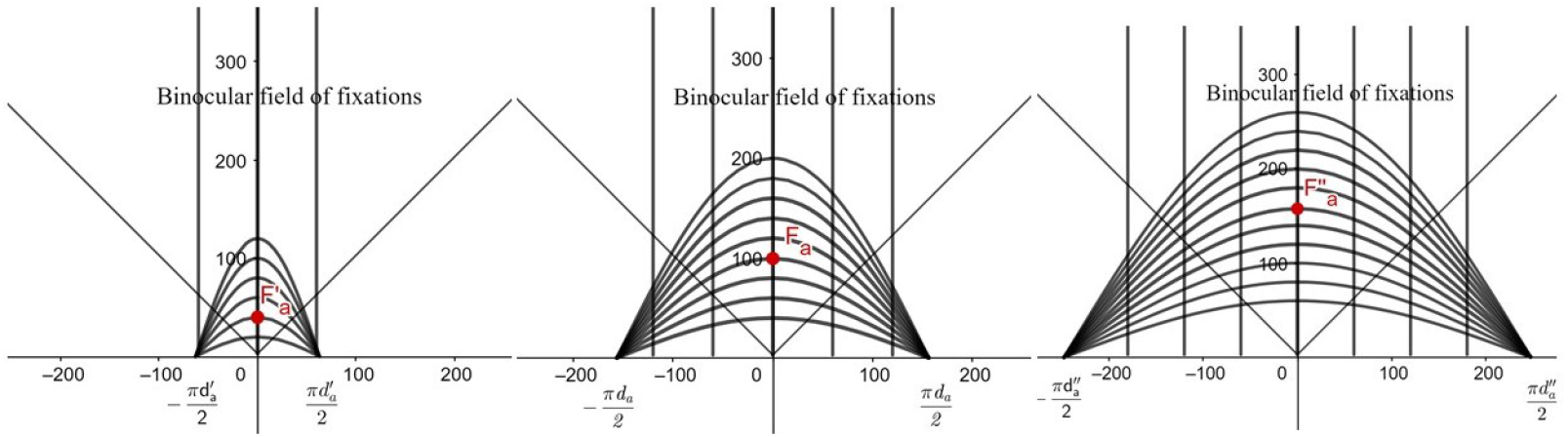
The geodesics in the resting eyes visual space for (a) *d*_*a*_ = 40.2 cm, (b) *d*_*a*_ = 99.61 cm, and (c) *d*_*a*_ = 157.75 cm. The curved geodesics start in *∂*_*y*_ direction from the points (0, *z*_0_). They are extended to the whole interval (−∞,∞) and approach asymptotically two points on the *y*-axis at ± (*π/*2)*d*_*a*_. The vertical geodesics start in *∂*_*z*_ direction from the points (*y*_0_, *ξ*). They are defined on the maximal interval 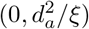.

The geodesics in the gaze direction *∂*_*z*_, that is, geodesics (38), can be analysed in details. Solving *z*_2_(*t*_2_) = *z*_2_ for *t*_2_, I derive the distance that does not depends on *y*_0_,

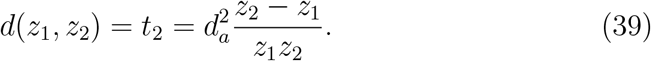

The total length of the geodesic from the point (0, *ξ*), the closest point that can be fixated binocularly, is

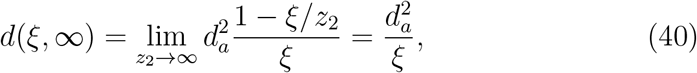

meaning that the *z*-geodesic has the finite length 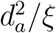 and, therefore, defines the visual horizon. This geodesic is incomplete.

Further, the distance from (0, *ξ*) to the eyes’ resting posture fixation is 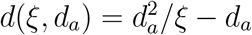, which means that the distance from the horopter to the horizon is *d*_*a*_.

### 8.1. Geodesic Curvature of Iso-disparity Curves

The iso-disparity curves in physical space are Euclidean lines of constant *z*-values. By the results of the previous section, they are not geodesics. For example, the Euclidean line through *F*_*a*_ differs from the geodesic line passing through the same point that is shown in Figure 8. How far this iso-disparity line differs from the geodesic is given by its geodesic curvature since every geodesic has zero geodesic curvature, see [22], page 137.

The unit speed iso-disparity curve is (*y*(*t*), *z*_0_) = (*z*_0_*/d*_*a*_)*t, z*_0_). It has the acceleration components

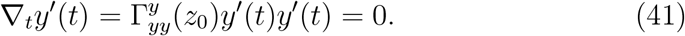

and

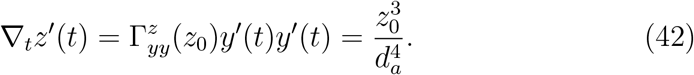

The geodesic curvature of the iso-disparity curve is

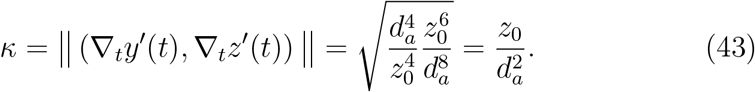

Thus, the iso-disparity line (a Euclidean line) for the resting eyes at the abathic distance *d*_*a*_ that is passing through (0, *z*_0_) is a curve with a constant geodesic curvature 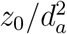. In particular, the horopteric line, in this case, has the geodesic curvature 1*/d*_*a*_.

## 9. The Riemann Curvature

Here the derivation of the Riemann curvature of visual space is only sketched for the eyes’ resting vergence posture and the reader is referred to Appendix Appendix A.4 for a more complete introduction to this complicated object. In the basis vectors *∂*_*i*_, where the index runs here over *y* and *z*, the covariant differentiation ∇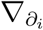 on the basis vectors is

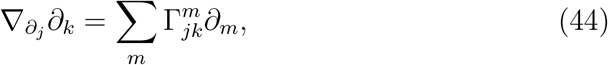

Following [22], the index notation of the curvature tensor can be written as,

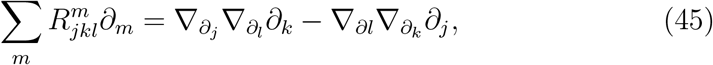

which, in low indices, is the following:

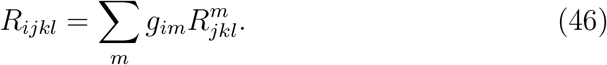

Using the notation corresponding to coordinates (*y, z*): (15); (16); (17); (18) and (44), the Riemann curvature tensor can be calculated based on (A.18) as follows:

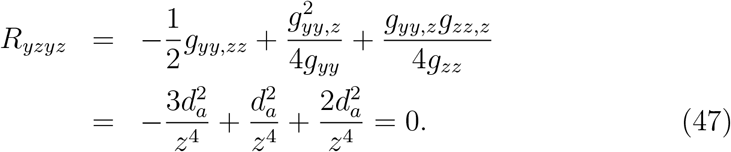

Thus, in this case of the eyes’ resting vergence posture, the curvature tensor of visual space vanishes identically and visual space is flat for this particular eye fixation [22].

## 10. Discussion

We note first, as the simulations clearly show, that visual space geometry is shaped by physical space being expanded between the head and the frontal line horopter at the abathic distance *d*_*a*_ and compressed beyond the horopter. These changes in perceived depth when the eyes are in their resting vergence posture (corresponding to the eye’s lens tilt’s angle *β*) make the geodesics incomplete such that visual space is bounded by the horizon in the direction of the gaze.

For the numerical calculations, I assume the human average lens tilt *β* = 3.3^*°*^, which gives the average abathic distance *d*_*a*_ = 99.61 cm [4]. I also estimate that *ξ* = 3 cm. Using these values in the geodesic distance expressions, I obtain the total distance to the horizon,

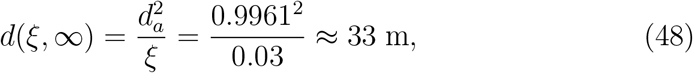

and the distance to the horopter at the abathic distance fixation,

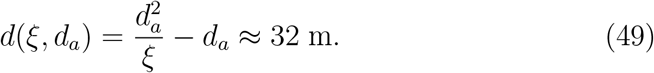

Therefore, the distance from the horopter at the abathic distance to the horizon is *d*(*d*_*a*_, ∞)≈1 m.

The finite distance to the horizon should be expected as we perceive objects in the horizontal binocular field against the horizon with the depth perception resulting from the metric (10) such that our field of vision appears two-dimensional with the objects initially located in visual processing at different distances from the horopter and finally perceived against the egocentric coordinates.

In literature, the distance of the vanishing points at the horizon is reported to be between 6 m and 100 m, see [23]. Here, the AE model in the binocular system determines the eyes’ resting vergence posture abathic distance that depends on the lens tilt angle *β*. The simulations were carried out for three values of the abathic distances; *d*_*a*_(0.5^*°*^) = 40.2 cm, *d*_*a*_(3.3^*°*^) = 99.61 cm, and *d*_*a*_(4^*°*^) = 157.75 cm. For these values, I obtain the distance to the horizon 5.4 m, 33 m, and 83 m, respectively. Thus, the geometric theory of the binocular perception developed in [4] and extended here to iso-disparity curves, unifies in one theory the distances to the horizon reported in the literature, where precise explanations for the differences in these distances were not provided.

The curvature of visual space, derived here for the eyes’ resting vergence posture when they fixate at the abathic distance, is zero, so the visual space is flat for this particular eyes posture. In this case, the iso-disparity curves in physical space are *z*-coordinates. However, the iso-disparity curve through (0, *z*_0_) for fixation at the abathic distance *d*_*a*_ has a non-zero geodesic curvatures 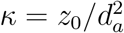. The geodesics starting at (0, *z*_0_) in direction *∂*_*y*_ consists families such that for each *d*_*a*_, the geodesics asymptotically converges to two points ± ((*π/*2)*d*_*a*_, 0) on the *y*-axis, see Figure 8. Therefore, they are parallel.

For all other fixations, the iso-disparity conics consist of the two families of iso-disparity curves in Figure 4 and Figure 5: for short-distance fixations the iso-disparity curves are ellipses, while for far-distance fixations the iso-disparity curves are hyperbolas. For the precise classification of the horopter’s shape and, hence, the iso-disparity curves’ shape with respect to the fixation point in the binocular field, see [4].

Because the convergence in the case of the elliptic iso-disparity family, the metric tensor depends on the *y* coordinates such that the geodesics in the Cyclopean gaze direction are not straight lines and the curvature of visual space should change from zero to positive curvature value. Similarly, because of the divergence in the case of the hyperbolic iso-disparity family, the curvature of visual space should be negative. The numerical value of the curvature cannot be readily obtained without the known metric tensor [24].

The results obtained here, although not complete, revise the abstract geometric theories of binocular phenomenal space, most notably, a constant negative-curvature hyperbolic geometry of visual space originally proposed by Luneburg [25] in the 1940s and further developed in [26] and [27], among other publications.

Luneburg’s approach is one of three theories that used axiom-based model geometries: Euclidean, hyperbolic, and spherical, to model phenomenal spatial relations [28]. These approaches, however, were not supported by experimental data and did not pay attention to biological and psychological aspects of human vision. For example, Luneburg’s theory uses the Vieth-Müller circle (VMC) as the geometric horopter. In [29], I showed that the VMC is anatomically inaccurate as a geometric horopter. In fact, it incorrectly assumes the nodal point located at the eye’s rotation center [30], which provides a good numerical approximation of the geometric horopter close to the fixation point. More significantly, when the VMC is compared with the simulated horopteric hyperbolas in [4], their difference in periphery is very large. Thus, the modeling based on the VMC will significantly distort the ambient space geometry provided by the coarse disparity, see the discussion in Section 5.

Beyond these three theories, there are also theories based on Riemann’s approach to studying abstract space with defined metric function. One of the most general distance-defined spaces is called metric space. These theories model visual space’s measurable properties such as distances and angles while also including stimulus location in physical space and psychological factors [28]. The theory proposed here for modeling visual space spatial relations is similar to the metric-based theories but also includes anatomical and physiological aspects of the human visual system.

Of the many other studies reporting results on phenomenal geometry, the experimentally obtained values of the visual space’s curvature in [31] could be closest to the results obtained in the presented here studies.

[31] found that the curvature changes from elliptic in near space to hyperbolic in far space. For very large distances, the visual plane became parabolic. Although, the values of the curvature can only be conjecture from Figure 9, in general it support the experimental results in [31].

**Figure 9:**
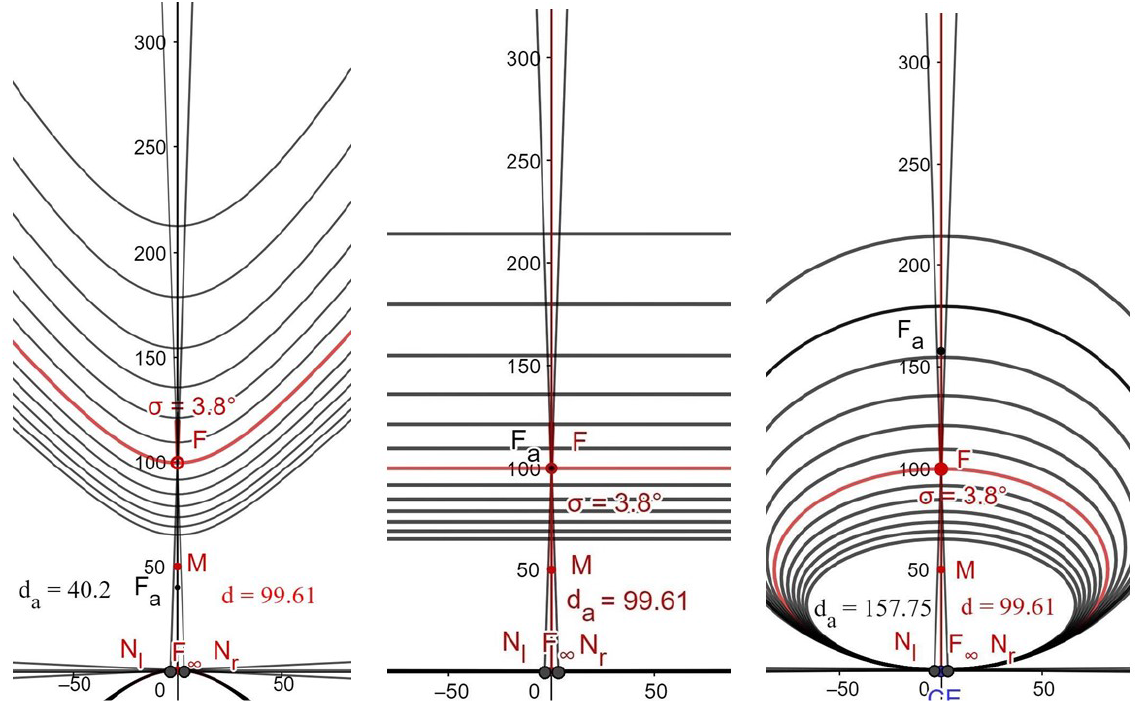
Three simulations for the same fixation and different values of the AE’s lens’ tilt. On the left, the lens’ tilt *β* = 0.5^*°*^, in the middle, *β* = 3, 3^*°*^, and on the right, *β* = 4^*°*^. In these cases, the curvature should be negative, zero and positive, respectively.

## 11. Closing Remarks

In this study, the geometries of visual space are discussed in terms of the iso-disparity curves constructed in the binocular system with the asymmetric eyes and simulated in *Geogebra*’s dynamic geometry software. In the eyes resting vergence posture, that is, when the eyes are fixated at the abathic distance, which depends on the eye’s asymmetry parameters, the iso-disparity curves are frontal straight lines. Otherwise, for all different fixations, the iso-disparity curves consist of ellipses or hyperbolas. They generally are perceived in the gaze direction as frontal lines.

Numerical analysis of the iso-disparity lines for the resting eyes, suggested the choice of the metric in visual space for this particular eyes’ posture. For a stationary, upright head, in the head coordinate system (*y, z*), where *y*-axis passes through the eye rotation centers and the *z*-axis is in the midsagittal direction, the metric is given by

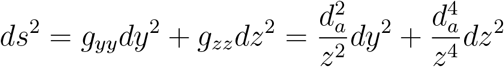

where *d*_*a*_ is the abathic distance. The distance in the gaze direction is

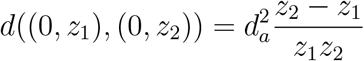

such that the total distance is 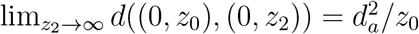 where (0, *z*_0_) is the closest point on which eyes can fixate. Thus, the Riemannian geometry framework furnishes, in this case, incomplete geodesics in the gaze direction giving a finite distance to the horizon. Also the curvature, in this case, is zero. Further, the iso-disparity line through (0, *z*_1_) has the geodesic curvature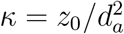, meaning that it is not a geodesic.

Because I postulate metric in this study, I briefly discuss three closely related metrics. Two of them have zero curvature and, in both cases, the iso-disparity lines are geodesics. The first of the metrics,

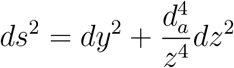

gives the same finite distance to the horizon. The second metric

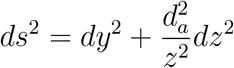

gives the distance *d*((0, *z*_1_), (0.*z*_2_)) = *d*_*a*_(ln *z*_2_ − ln *z*_1_) and, hence, the horizon infinitely away.

The third closely related metric,

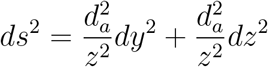

gives the Riemann curvature, 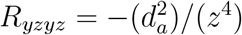, and, therefore, the Gaussian curvature

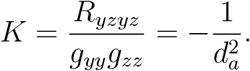

The above discussion shows that when the iso-disparity curves are frontal straight lines and the curvature is zero, the geodesics in the binocular gaze direction are straight lines, possibly incomplete.

## APPENDIX

### Appendix A. Riemannian Geometry Primer

The binocular space of fixations, *B*, is a convex subset in the horizontal visual plane discussed in Section 3. The global coordinate system (*x*^*i*^) is specified relative to the stationary upright head where the index *i* runs over *y* and *z*. The coordinates (*x*^*i*^) give the basis vectors *∂*_*i*_ introduced in Section 6. The metric tensor *g*(*∂*_*i*_, *∂*_*j*_) = *g*_*ij*_ on *B* is obtained from the geometric analysis of disparity-based perception. It define the length of the tangent vector and the angle between two tangent vectors as discussed in Section 6. To avoid technicalities, I review here the basic notions of Riemannian geometry in one global chart. Also I use so called *Einstein summation convention*: when an index occurs twice in the same expression in upper and in lower positions, then the expression is implicitly summed over all possible values for that index.

#### Appendix A.1. Tangent Vectors and Metric

Let *φ* : *M* → *R*^*n*^, *φ*(*p*) = (*x*^*i*^), be a global chart on a manifold *M* of dimension *n*. The tangent vector *X*_*p*_ at *p* ∈ *M* is identified with the directional derivative on *f* ∈ *C*^*∞*^(*M*) viz.

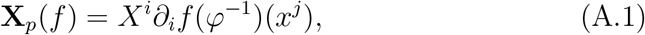

where *∂*_*i*_ = *∂/∂x*^*i*^ are partial derivatives. Thus, this tangent vector at *p*, is **X**_*p*_ = *X*^*i*^*∂*_*i*_ | _(*p*)_. The set (*∂*_1_, *∂*_2_, …, *∂*_*n*_) gives vector basis of the tangent space *T*_*p*_*M* of *M* at *p*. The tangent space *T*_*p*_*M* is identified with *R*^*n*^ that has the origin at *p*. The tangent vector field on *M* is a map **X** on *M* such that **X**(*p*) = **X**_*p*_.

*Riemannian space* is a manifold *M* equipped with a *Riemannian metric g* that for each *p* ∈ *M* defines the scalar product *g*_*p*_ in the tangent space *T*_*p*_*M* that depends smoothly on *p*. For any vector **X** ∈ *T*_*p*_*M, g*_*p*_(**X, X**) defines the square of the length of **X**. In the basis *∂*_*i*_ in *T*_*p*_*M, g*_*p*_(*∂*_*i*_, *∂*_*j*_) = *g*_*ij*_. Then, for any two vectors **X** = *X*^*i*^*∂*_*i*_ and **Y** = *Y* ^*i*^*∂*_*i*_ in *T*_*p*_*M, g*_*p*_(**X, Y**) = *g*_*ij*_*X*^*i*^*Y* ^*j*^.

#### Appendix A.2. Covariant Derivative and Connection

For a given tangent vector field **X** = *X*^*i*^*∂*_*i*_, the covariant derivative ∇_**X**_ on a function *f* on *M* is defined as the directional derivative

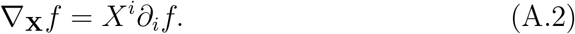

To obtain the covariant derivative of a vector field **Y** = *Y* ^*i*^*∂*_*i*_ along **X** = *X*^*i*^*∂*_*i*_, the previous differentiation is extended as follows

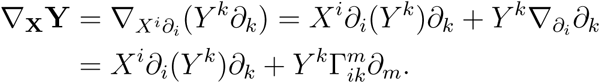

where in the last equality the vector field 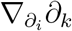 is decomposed in the basis *∂*_*i*_:

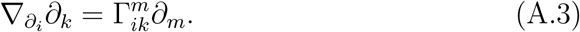

The coefficients 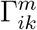 are known as Christoffel symbols in the coordinates (*x*^*i*^).

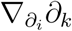 can be interpreted as comparison of vectors *∂*_*k*_ in infinitesimally close tangent spaces in the direction of *∂*_*i*_. Thus, the covariant derivative establishes *connection* ∇ between tangent spaces, cf. the discussion in Section 6. Recall that the formulation here is global such that I do not discuss the local aspects of the covariant derivative.

A connection ∇ is symmetric if

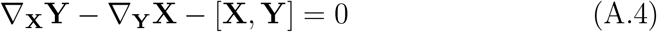

where [**X, Y**] is the commutator: [**X, Y**]*f* = **XY***f* − **YX***f*. Because in the basis *∂*_*i*_, [*∂*_*i*_, *∂*_*j*_] = 0, then we have

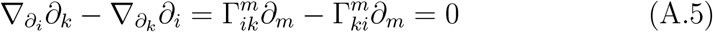

and, therefore, Christoffel symbols are symmetric 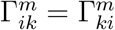 in lower indices.

A symmetric connection is *Levi-Civita connection* if it preserves the metric: for any two vector fields **Y, Z**,

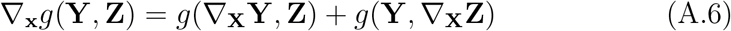

I state the following theorem without proving it.

##### Theorem.

*There exists unique Levi-Civita connection on the Riemannian manifold* (*M, g*). *In local coordinates Christoffel symbols of Levi-Civita connection are given by*

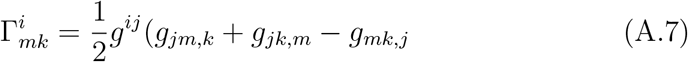

*where* (*g*_*ij*_) *is the Riemannian metric and* (*g*^*ij*^) *is the inverse of* (*g*_*ij*_).

#### Appendix A.3. Parallel Transport and Geodesics

Let **X**(*t*) be a vector field along a curve *C* : *x*(*t*) = (*x*^*i*^(*t*)), *t*_0_≤*t*≤*t*_1_ and **v** = *dx*(*t/dt* = *dx*^*i*^(*t*)*/dt∂*_*I*_ | _*x*(*t*)_ the velocity vector of *C*. We say that **X**(*t*) is a parallel transport of **X** = **X**(*t*_0_) along *C* if the covariant derivative along *C* is zero

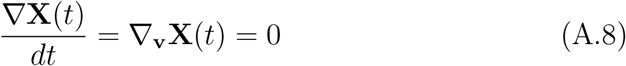

In coordinates, it can be written as

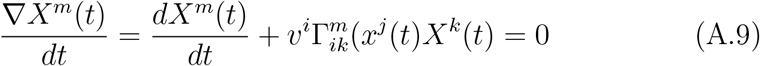

Because ∇ is Levi-Civita connection, it is easy to show that the Riemannian metric is *preserved* during the parallel transport.

A parametrized curve *C* : *x*^*i*^(*t*) is geodesic if parallel transport of velocity vector *v*^*i*^(*t*) = *dx*^*i*^(*t*)*/dt* along the curve *C* is a velocity vector at any point of *C*, that is, if

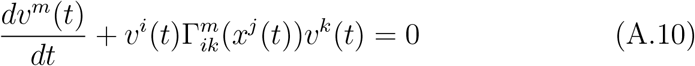

i.e.,

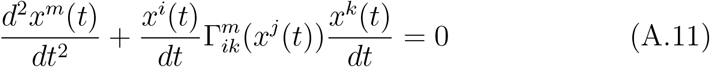

Because Levi-Civita connection preserves the Riemannian metric, the length of velocity vector is constant along the geodesic. The geodesic that has unit speed is parametrized by the geodesic’s arc length.

#### Appendix A.4. Riemann Curvature Tensor

Coordinate-invariant definition of the Riemann curvature tensor for Levi-Civita connection ∇ is given as

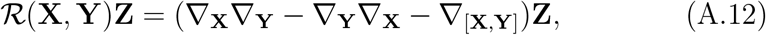

where **X, Y** and **Z** are vector fields. It is a complicated object and I introduce it briefly. In coordinates, the components of the curvature tensor are obtained as follows

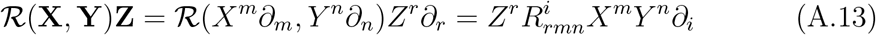

where

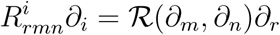

Following [22], using that [*∂*_*i*_, *∂*_*j*_] = 0, in the vector basis *∂*_*i*_ the index notation of the curvature tensor is given by

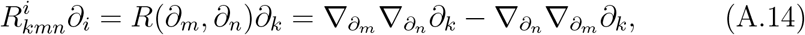

which, by substituting 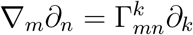 can be written as

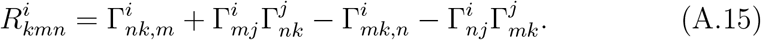

in terms of the Christoffel symbols. Then, the Riemann curvature tensor with low indices is

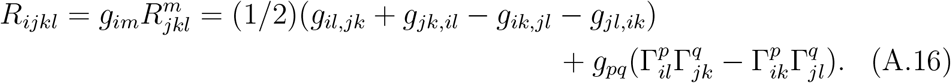

Now, the curvature tensor for Levi-Civita connection obeys the following identities

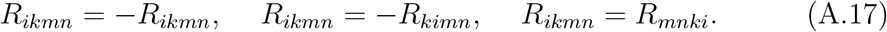

For two-dimensional Riemannian manifolds these identities imply that *R*_1212_ yields all components up to a sign. Thus,

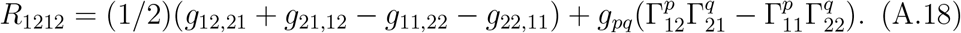

## Acknowledgment

I thank Alice Turski for her help in editing this manuscript.

## References

[1] R. A. Armstrong, R. C. Cubbidge, The eye and vision: an overview, in: Handbook of Nutrition, Diet, and the Eye, Academic Press, San Diego, 2019, pp. 3–14. doi:10.1016/B978-0-12-815245-4.00001-6.

[2] R. W. Guillery, C. A. Mason, J. S. H. Taylor, Developmental determinants of the mammalian optic chiasm, J. Neurosci. 15 (1995) 4727–4737. doi: 10.1523/JNEUROSCI.15-07-04727.

[3] K. N. Ogle, Precision and validity of stereoscopic depth perception from double image, J. Opt. Soc. Am. 41 (1953) 906–913.

[4] J. Turski, A geometric theory integrating human binocular vision with eye movement, Front. Neurosci. 14:555965 (2020) 1–17. doi:10.3389/fnins.2020.555965.

[5] J. Turski, Binocular system with asymmetric eyes, J. Opt. Soc. Am. A 35 (2018) 1180–1191. doi:10.1364/JOSAA.35.001180.

[6] P. Artal, Optics of the eyes and its impact in vision, Adv. Opt. Photon 6 (2014) 340–367. doi:10.1364/AOP.6.000340.

[7] K. Ogle, Researches in Binocular Vision, PA:WB Saunders, Philadelphia, 1950.

[8] T. Shipley, S. Rawlings, The nonius horopter-i. history and theory, Vis. Res. 10 (1970) 1225–1262. doi:10.1016/0042-6989(70)90039-8.

[9] G. von Noorden, E. C. Campos, Binocular Vision and Ocular Motility: Theory and Management of Strabismus, Mosby, A Harcourt Health Sciences Co., St. Louis., 2002.

[10] T. Shipley, S. Rawlings, The nonius horopter–ii. an experimental report, Vis. Res. 10 (1970) 1263–1299. doi:10.1016/0042-6989(70)90040-4.

[11] W. Jaschinski-Kruza, Eyestrain in vdu users: viewing distance and the resting position of ocular muscles, Hum. Factors 33 (1991) 69–83. doi:10.1177/001872089103300106.

[12] S. M. Ebenholtz, Oculomotor Systems and Perception, Cambridge U. Press, 2008. doi:10.1017/CBO9780511529795.

[13] D. Guitton, M. Volle, Gaze control in humans: eye-head coordination during orienting movements to targets within and beyond the oculomotor range, J. Neurophysiol. 58 (1987) 427–459. doi:10.1152/jn.1987.58.3.427.

[14] G. Maiello, M. Chessa, P. J. Bex, F. Solari, Near optimal combination of disparity across a log-polar scaled visual field, PLoS Comp. Biol. 16 (2020) e1007699. doi:10.1371/journal.pcbi.1007699.

[15] A. M. Norcia, E. E. Sutter, C. W. Tyler, Electrophysiological evidence for the existence of coarse and fine disparity mechanisms in human, Vis. Res. 25 (1985) 1603–1611. doi:10.1016/0042-6989(85)90130-0.

[16] C. Blakemore, The range and scope of binocular depth discrimination in man, J. Physiol. 211 (1970) 599–622.

[17] L. M. Wilcox, R. S. Allison, Coarse-fine dichotomies in human stereopsis, Vis. Res. 49 (2009) 2653–2665. doi:10.1016/j.visres.2009.06.004.

[18] S. R. Barry, Beyond the critical period. acquiring stereopsis in adulthood, in: Plasticity in Sensory Systems, Cambridge U. Press, Cambridge, 2013, pp. 175–195. doi:10.1017/CBO9781139136907.010.

[19] M. P. Do Carmo, Riemannian Geometry, Birkhäuser, Boston, 1993.

[20] V. P. Ermakov, Second order differential equations. conditions of complete integrability, Universita Izvestia Kiev 9 (1880) 1–25. (Russian).

[21] E. Pinney, The nonlinear differential equation y″ + p(x)y′ + cy-3, Proc. Am. Math. Sc. 1 (1950) 681.

[22] J. M. Lee, Riemannian Manifolds: An Introduction to Curvature, Springer, NY, 1997.

[23] C. J. Erkelens, Perspective space as a model for distance and size perception, i-Perception (2017) 1–20doi:10.1177/2041669517735541.

[24] K. Burns, V. S. Matveev, Open problems and questions about geodesics, Ergodic Theory and Dynamical Systems 41 (2021) 641–684.

[25] R. K. Luneburg, Mathematical Analysis of Binocular Vision, Princeton U. Press, USA, 1947.

[26] A. A. Blank, The luneburg theory of binocular visual space, J. Opt. Soc. Am. 43 (1953) 717–727.

[27] T. Indow, A critical review of luneburg’s model with regard to global structure of visual space, Psychological Review 98 (1991) 430–453. doi: 0033-295X.98.3.430.

[28] M. Wagner, The Geometries of Visual Space, Lawrence Erlbaum Associates, Mahwah, NJ, 2005.

[29] J. Turski, On binocular vision: the geometric horopter and cyclopean eye, Vis. Res. 119 (2016) 73–81. doi:10.1016/j.visres.2015.11.001.

[30] W. L. Gulick, R. B. Lawson, Human stereopsis: A Psychophysical Analysis, Oxford U. Press, NY, 1976.

[31] J. J. Koenderink, A. J. van Doom, J. S. Lappin, Direct measurement of the curvature of visual space, Perception 29 (2000) 69–79. doi: 10.1068/p2921.

